# A non-canonical arm of UPR^ER^ mediates longevity through ER remodeling and lipophagy

**DOI:** 10.1101/471177

**Authors:** Joseph R. Daniele, Ryo Higuchi-Sanabria, Vidhya Ramachandran, Melissa Sanchez, Jenni Durieux, Sarah U. Tronnes, Joseph W. Paul, Daniel J. Esping, Samira Monshietehadi, Melissa G. Metcalf, Andrew Dillin

## Abstract

Longevity is dictated by a combination of environmental and genetic factors. One of the key mechanisms implicated in regulating lifespan extension is the ability to induce protein chaperones to promote protein homeostasis. However, it is unclear whether protein chaperones exclusively regulate longevity. Previous work has shown that activating the unfolded protein response of the endoplasmic reticulum (UPR^ER^) in neurons can signal peripheral tissues to promote chaperone expression, thus enhancing organismal stress resistance and extending lifespan. Here, we find that this activation not only promotes chaperones, but facilitates a dramatic restructuring of ER morphology in intestinal cells. This restructuring, which includes depletion of lipid droplets, ER expansion, and ER tubulation, depends of lipophagy. Surprisingly, we find that lipophagy is required for lifespan extension and is completely independent of chaperone function. Therefore, UPR induction in neurons triggers two distinct programs in the periphery: the canonical arm through protein chaperones, and a non-canonical mechanism through lipid depletion. In summary, our study identifies lipophagy as an integral component of UPR^ER^-induced longevity.

## MAIN

Life presents a myriad of challenges, stressors, and environmental shifts to which cells must adapt in order to survive and thrive. The homeostatic regulation of protein folding (proteostasis), which is monitored in specific subcellular compartments (the endoplasmic reticulum (ER), mitochondria, cytosol) is an integral player in stress resistance and longevity. The ER, in particular, is a central regulator of stress monitoring since it controls (1) nearly a third of the cell’s proteins, (2) provides an internal medium and transponder for lipid homeostasis and cell signaling, and (3) communicates directly with all other organelles to maintain cellular secretion.

Notably, the ER has evolved three primary branches of its unfolded protein response (UPR^ER^) to maintain proper secretion, protein folding, and lipid homeostasis. The most studied of the three UPR^ER^ branches involves the membrane localized/endoribonuclease inositol-requiring protein-1 (IRE-1), which upon unfolded protein stress in the ER lumen (or lipid disequilibrium in the ER membrane), will splice a specific intron from the mRNA of the transcription factor, X-Box binding protein-1 (xBP-1), to create *xbp-1s* which then upregulates the expression of protein degradation, protein folding, and lipid metabolism gene targets. Another branch of the UPR^ER^ is the membrane-localized protein kinase, R (PKR)-like endoplasmic reticulum kinase (PERK), which upon ER stress, globally reduces protein translation via phosphorylation of the translation initiation factor, eIF2α. Transcripts which promote protein folding and apoptosis, however, are preferentially translated. Finally, the last of the three UPR^ER^ branches is the ER-membrane-localized transcription factor-6 (ATF6), which upon luminal stress, moves to the Golgi to become a mature transcription factor capable of upregulating canonical proteostasis genes (Frakes and Dillin, 2017; Walter and Ron, 2011).

While these pathways have been intensively studied for the last two decades, much less is known about the adaptive responses of the ER under long-lived conditions. The small nematode, *C. elegans,* is an excellent model to simultaneously study the genetic underpinnings of cell-cell communication under adaptive conditions, while also enabling the monitoring of its cell biology and organismal health (e.g. lifespan, stress resistance). Work with *C. elegans* has shown that its cells become less capable in protein folding and also less able to induce stress responses to proteotoxicity with advanced age (Ben-Zvi et al., 2009; Brown and Naidoo, 2012; Dillin et al., 2002a, 2002b; Durieux et al., 2011; Higuchi-Sanabria et al., 2018a; Taylor and Dillin, 2013). Interestingly, overexpression (OE) of the spliced version of *xbp-1 (xbp-1-s)* specifically in neurons, extends organismal lifespan and increases ER stress tolerance in a cell non-autonomous manner (Taylor and Dillin, 2013). While the precise, small ER-stress signal (SERSS) was not identified, small clear vesicles (SCV) are required for this beneficial effect, which could be host to numerous neurotransmitters. The intestine, the putative target tissue in this paradigm, induces chaperones in an XBP-1-dependent manner. However, it is well understood that UPR^ER^ induction does not alter only protein homeostasis through chaperones, but is an important mechanism for metabolic change and handling of lipid stress (Shen et al., 2005).

We hypothesized that induction of the UPR^ER^ in neurons, which reverses the age-dependent loss of ER proteostasis, also enacts a dramatic restructuring of ER morphology which, in turn, imparts a beneficial metabolic change and promotes longevity. Although whole-organismal metabolic restructuring has been a topic of intense study in the aging field (e.g. Finkel, 2015), much less is known about the adaptive responses of organelles (e.g. the ER) in long-lived conditions. Here, we find that upon neuronal *xbp-1s* OE, peripheral, non-neuronal cells in this paradigm possess distinct ER structures, which we call “circular ER-derived membranes” (CERMs), and that this morphology is dependent on the lipophagic depletion of intestinal lipid droplets (LDs). The restructuring and lipid depletion mediated by a conserved protein complex (RAB-10/EHBP-1/RME-1), which is necessary for the metabolic change and increased longevity seen in neuronal *xbp-1s* animals. Perhaps most interesting is that changes in ER remodeling and lipid depletion is uniquely distinct from the canonical UPR^ER^ involving chaperone induction, such that perturbations in lipid depletion abrogates the beneficial effect of UPR^ER^ induction, despite not altering the induction of chaperones. Thus, we argue that the beneficial effects of non-autonomous UPR^ER^ is dependent on two independent, yet equally important, arms of UPR^ER^: the canonical arm including chaperone induction to mediate protein homeostasis and the the non-canonical arm, which induces ER remodeling and lipophagy.

### Ectopic UPR^ER^ induction results in ER remodeling and lipid depletion

Although it is assumed that ER integrity and function breaks down with age, relatively little is understood about the mechanisms governing this process, especially in organismal models of extended longevity (Gordon, 1971; Minakshi et al., 2017; Palay and Palade, 1955). Here, we visualized the ER during various stages of adulthood in *C. elegans* using an mCherry::HDEL fusion protein that localizes to the ER lumen. At Day 1 of adulthood, the ER fills the intestinal lumen and can be visualized as a sponge-like, interconnected, and uniform structure (**Fig. 1a, top**). As early as Day 4, intestinal ER is less uniform and begins to collapse, forming disorganized, aggregated structures, with increased severity by Day 7.

**Figure 1.**
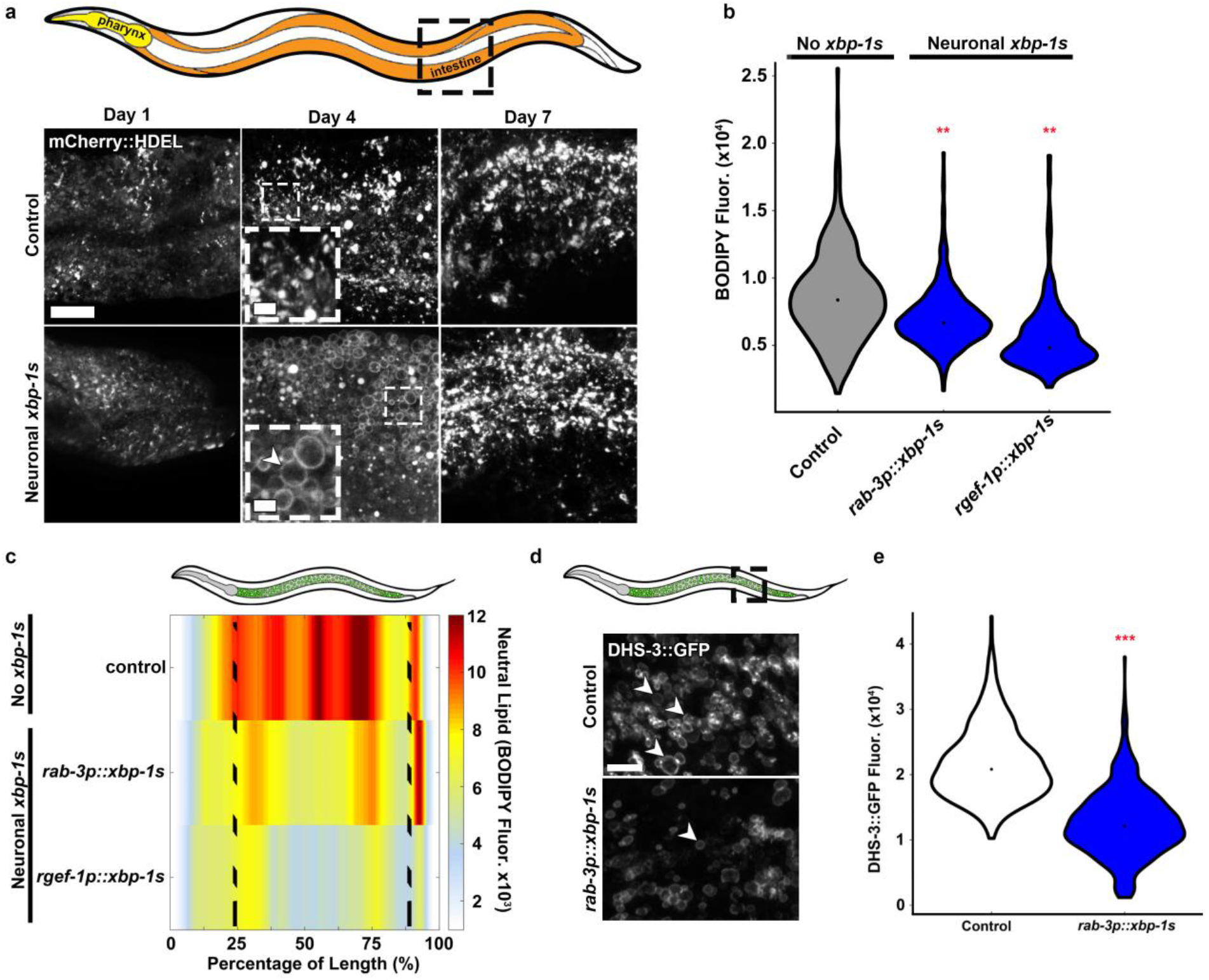
Cell non-autonomous UPR^ER^ remodels the cell biology and metabolism of the target tissue. **(a)** Representative micrographs of endoplasmic reticulum (ER) morphology (via an intestine-specific *vha-6p::mCherry::HDEL)* at various times in adulthood (early Day 1, Day 4, and Day 7) under “control” and cell non-autonomous UPR^ER^ activated (“Neuronal *xbp-1s”, rab-3p::xbp-1s)* conditions. In cartoon, intestine is shown in orange and pharynx in yellow. Arrowhead indicates representative CERM. Scale = 10 μm. **(b)** Quantification of neutral lipid amount (via BODIPY 409/503 staining) of whole nematodes upon expression of *xbp-1s* in neurons using two different promoters. Grey plots indicate co-injection markers alone (controls) and blue plots indicate neuronal *xbp-1s* OE. \*\**P*≤0.01 against all controls. **(c)** Median profiles of nematode neutral lipid distribution (via BODIPY 409/503 stain) in b following alignment. Data is shown as a heat map with Red as the highest amount of lipid and white the lowest. All animals are positioned with the anterior side to the left. Numbered lines depict independent extrachromosomal arrays with or without *xbp-1s* overexpression constructs. **(d)** Representative micrographs of intestinal lipid droplets (LDs) (via *dhs-3p::DHS-3::GFP*) in Day 2 adult control and neuronal *xbp-1s* animals. Scale = 5 μm. Arrowheads indicate representative LDs. **(e)** Quantification of lipid droplets (via DHS-3::GFP fluorescence) in Day 2 adults in control (white) and neuronal *xbp-1s* (blue) animals. \*\*\**P*<0.001, by Mann-Whitney, against age-matched “Control”. Plots are representative of three biological replicates and derived from *n* ≥ 50 organisms for each condition.

To determine whether preserving ER protein homeostasis would prevent the age-associated decline in ER function, we interrogated ER structure and organization in a long-lived model of *C. elegans*, which maintains proper ER protein homeostasis late into adulthood. Here, UPR^ER^ is activated specifically in neurons by ectopic overexpression of *xbp-1s* using a neuron-specific promoter to bypass the need for stress-induced, IRE-1-mediated cleavage of *xbp-1.* These animals, hereon referred to as neuronal *xbp-1s* animals, have constitutive UPR^ER^ induction in neurons, which can signal non-autonomously to peripheral tissues and activate UPR^ER^, ultimately resulting in lifespan extension (Taylor and Dillin, 2013). Neuronal *xbp-1s* animals display no difference in intestinal ER structures at Day 1 compared to wild type animals. Unexpectedly, at Day 4 when wild-type animals show collapse of the ER, neuronal *xbp-1s* animals have a spherical ER morphology that is not seen at any age in wild type animals (**Fig. 1a, bottom, see arrowhead**). By Day 7, these spherical structures are gone and the ER network collapses in a manner indistinguishable from wild type animals. Therefore, constitutive activation of UPR^ER^ is not sufficient to protect ER quality and structure during aging, and only show significant differences in early adulthood. For ease of clarification, we termed theses transient ER structures <u>c</u>ircular <u>ER</u>-derived <u>m</u>embranes (CERMs), which could be observed as early as Day 2 of adulthood in neuronal *xbp-1s* animals (**Fig. 1a, see arrowheads**). This is not specific to a single neuronal promoter, as overexpression of *xbp-1s* using a different neuronal promoter, *rgef-1p,* resulted in similar CERM formation and lifespan extension (**Extended Figure 1a-b**). A lack of cytosol inside these structures indicates they are discrete and not solely a result of increased ER tubulation (**Extended Data Fig.1c**).

To determine whether CERM formation was unique to the neuronal, non-autonomous, paradigm of UPR^ER^ activation, we tested whether systemic-or tissue-specific (muscle or intestine) overexpression of *xbp-1s* resulted in formation of intestinal CERMs. Interestingly, overexpression of *xbp-1s* in muscle or intestine did not result in CERM formation in the intestine (**Extended Data Fig. 1d**). These data suggest two potential models: 1) non-autonomous *xbp-1s* signaling from neurons results in a unique response distinct from autonomous UPR^ER^ activation by *xbp-1s* expression in the intestine or 2) there is a dosage effect of *xbp-1s* on CERM formation, and neuronal *xbp-1s* results in higher *xbp-1s* expression in intestine compared to *xbp-1s* driven by the intestinal promoter *(gly-19p)* alone. To our surprise, all-tissue overexpression of *xbp-1s* did not result in CERM formation, despite the presence of *xbp-1s* overexpression in the neurons in this strain (**Extended Data Fig. 1d**, compare neuronal *xbp-1s* to all tissue *xbp-1s).* These data provide further evidence for the requirement of a critical threshold of *xbp-1s* in neurons and/or intestine to promote CERM formation. Alternatively, it is possible that intestinal overexpression of *xbp-1s* perturbs non-autonomous signaling from neurons. Nonetheless, neuronal activation of the UPR^ER^ results in a dramatic restructuring of the ER that is not found in wild-type animals or other paradigms of UPR^ER^ activation studied here.

The appearance of CERMs only in neuronal *xbp-1s* animals suggests that perhaps the ER remodeling can be linked to the extreme longevity of these animals, as the other models of *xbp-1s* overexpression fail to increase lifespan to the same extent. To investigate the nature of the cell non-autonomous induction of CERMs, we first tested whether UPR^ER^ activation in the intestine was essential for CERM formation. Neuronal *xbp-1s* animals have UPR^ER^ induction in distal tissue, such as the intestine, and this distal activation of UPR^ER^ is essential for their prolonged lifespan (Taylor and Dillin, 2013). Specifically, knockdown of *xbp-1* in peripheral tissue through RNAi resulted in a loss of UPR^ER^ activation in intestine and ultimately a suppression of the lifespan extension in neuronal *xbp-1s* animals (Taylor and Dillin, 2013). Similarly, we find that CERM formation in neuronal *xbp-1s* animals is also dependent on *xbp-1* in distal tissues (**Extended Data Fig. 1e**). Therefore, distal induction of the UpR^ER^ and CERM formation by neuronal *xbp-1s* is dependent upon *xbp-1* in the periphery.

As a critical organelle in secretory vesicle formation, we hypothesized that the major ER remodeling found in neuronal *xbp-1s* animals could be due to increased intestinal secretion. To measure secretion, we compared VIT-2::GFP expression in neuronal *xbp-1s* animals compared to wild type. VIT-2::GFP, is a fluorescently tagged version of vitellogenin, a reporter for intestinal protein secretion (Grant and Hirsh, 1999; Stevens and Spang, 2013). This protein is integral to maternal yolk formation, and under normal conditions is secreted by the intestine and endocytosed by developing eggs. Defects in secretion can be visualized by increased intestinal fluorescence and decreased egg-localized fluorescence. Indeed, we find that neuronal *xbp-1s* animals have increased VIT-2::GFP expression in their eggs, suggesting a significant increase in intestinal secretion in these animals (**Extended Data Fig. 1f, see eggs in whole worms (top) and fluorescence of isolated eggs (bottom)**). A massive increase in secretion would result in lipid depletion due to the expansion of ER membranes. Thus, we next tested whether intestinal ER remodeling of neuronal *xbp-1s* animals occurred concurrently with lipid depletion. We measured intestinal lipids using a combination of whole-organism and tissue-specific quantification.

Our lab had previously developed a methodology and software, called LAMPro, to exploit underused positional information inherent to biosorter data, which takes fluorescent traces of many animals at once, and defines key regions of interest to various cellular phenomena (Daniele et al., 2017). This enabled us to overcome the practical limits of microscopy (e.g. low n) to reliably measure the distribution of total lipids across animals. Using LAMPro technology, we measured whole-body neutral lipid content (via BODIPY 409/503 staining). Briefly, animals were staged when CERMs were prominent, fluorescent integrated intensity was measured across the length of BODIPY-stained worms, and the data was represented as a heat map of median profiles of relative neutral lipid distribution. Notably, we found that neuronal *xbp-1s* animals show a dramatic depletion of lipids compared to wild type animals or animals overexpressing *xbp-1s* in muscle or intestine (**Fig. 1b-c, Extended Data Fig. 1g-h, compare grey plots to blue plots**).

Next, we tested whether the decline in lipid content is due to the loss of lipid droplets (LDs), the primary organelle that stores intestinal lipids in *C. elegans* (Lemieux and Ashrafi, 2015). Unlike most organelles, LDs are composed of a trigylceride and cholesteryl ester core surrounded by a phospholipid monolayer and decorated on the outside with proteins and sterol. We visualized LDs with the intestine-specific, fluorescently labeled, short-chain dehydrogenase, DHS-3::GFP fusion protein, which is abundant on the surface of *C. elegans* intestinal lipid droplets (Na et al., 2015; Zhang et al., 2012). Using this marker, we found that lipid droplet content was significantly lower in neuronal *xbp-1s* animals compared to wild-type animals (**Fig. 1d-e, see arrowheads**). This lipid depletion was only apparent in early adulthood (Day 1 to Day 4) in neuronal *xbp-1s* animals, but late-stage larval animals (L4) and Day 7 adult neuronal *xbp-1s* animals had similar lipid droplet content to wild type animals (**Extended Data Fig. 1i**). Moreover, this lipid depletion was dependent on peripheral *xbp-1* expression, as RNAi knockdown of *xbp-1* suppressed this phenotype, similar to CERM formation (**Extended Data Fig. 1j**). These data support the hypothesis that the ER-remodeling found in neuronal *xbp-1s* animals results in significant intestinal lipid depletion.

### Ectopic UPR^ER^ activation increases lysosomes in the intestine

Under conditions of UPR^ER^ activation and massive ER expansion, induction of ER-phagy and lipophagy can aid in the redistribution of lipids (Schuck et al., 2014; Vevea et al., 2015). For example, in yeast, ER stress causes ER expansion and the formation of ER whorls (defined as numerous and matching membranes organized in a regularly spaced, concentric manner), which are then targeted to the vacuole for selective degradation through autophagy (Bernales et al., 2006). Here, we questioned whether the ER remodeling and lipid depletion were due to ER-phagy or lipophagy during UPR^ER^ activation. Consistent with previous studies, we find that acute ER stress by treatment of animals with the chemical agent, tunicamycin (Tm), resulted in CERM formation, although at a much lower efficiency than neuronal *xbp-1s* overexpression (**Extended Data Fig. 2a**). More importantly, we find an increase in lysosomal number in neuronal *xbp-1s* animals, compared to wild-type animals, suggesting that the ER expansion and lipid depletion found in these animals is due to increased lipid turnover through autophagy (**Fig. 2a-b using LMP-1::GFP, 2c-d & Extended Data Fig. 2b using lysotracker**). Indeed, we see that intestinal CERMs found in neuronal *xbp-1s* animals colocalize with lysosomes with either lysotracker green or LMP-1, a lysosome-associated membrane protein (**Fig. 2a, Extended Data Fig. 2b**). Notably, lysosomal expansion in neuronal *xbp-1s* animals was dependent on peripheral *xbp-1* (**Fig. 2c-d, Extended Data Fig. 2b**).

**Figure 2.**
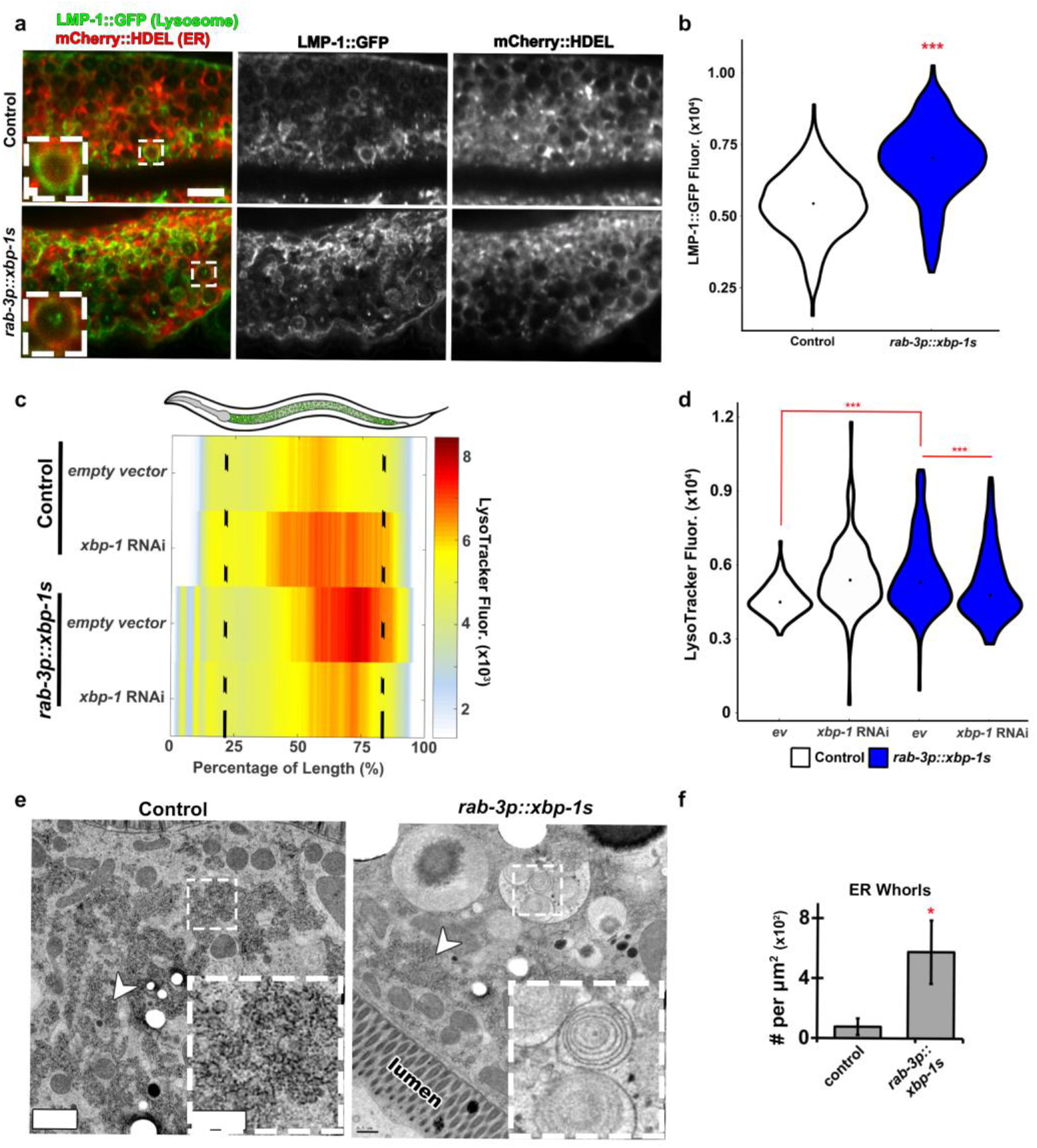
ER remodeling is coincident with more intestinal lysosomes. **(a)** Confocal micrographs of intestinal ER *(vha-6p::mCherry::HDEL)* and lysosomes *(vha-6p::lmp-1::GFP),* in fixed animals, with and without *rab-3p::xbp-1s.* Scale = 5 μm (and 0.5 μm for inset). **(b)** Quantification of lysosomes (via LMP-1::GFP) in control and *rab-3p::xbp-1s* animals. **(c)** Distribution of acidic lysosomes (via Lysotracker Green) in control (top) and *rab-3p::xbp-1s* (bottom) animals following profile alignments. Empty vector and *xbp-1* RNAi treatments are included. All animals are positioned with the anterior side to the left. **(d)** Total median quantification of lysosomes (via Lysotracker Green) in c. For b & d, \*\*\**P*<0.001 by Mann-Whitney. **(e)** Electron micrographs of intestine from control and neuronal *xbp-1s* overexpressing animals *(rab-3p::xbp-1s)*. Arrowheads mark rough ER. Scale = 1 μm (0.25 μm for inset). **(f)** Quantification of ER whorls per μm^2^ in control vs. *rab-3p::xbp-1s* animals. *P<0.05 by *t-test.* Plots are representative of three biological replicates and derived from *n* ≥ 50 organisms for each condition.

To further determine whether lysosomal expansion was specific to the neuronal *xbp-1s* paradigm, we tested whether wild type worms treated with exogenous ER stress also increased lysosomal content. Interestingly, worms treated with Tm do not show an increase in lysosomes, despite having signs of ER remodeling through CERM formation (**Extended Data Fig. 2a, compare control + Tm (bottom, left) to *rab-3p::xbp-1s* + DMSO (top, right) and see Extended Data Fig. 2c-d**). These data provide further evidence that the ER remodeling and lipid depletion phenomena found in neuronal *xbp-1s* animals is unique to non-autonomous signaling and not a general response to ER stress.

Finally, we sought to directly test whether neuronal *xbp-1s* resulted in increased lipophagy and/or ER-phagy. We measured the levels of a fluorescently-labeled lipophagy substrate, GFP::RAB-7 fusion protein, and detected less RAB-7 in these animals, suggesting that depleted lipids are likely due to an increase in lysosomal degradation (**Extended Data Fig. 2e**) (Guerra and Bucci, 2016; Schulze et al., 2017). Furthermore, high magnification transmission electron micrograph imaging of neuronal *xbp-1s* animals identified denser patches of rough ER compared to controls (**Fig. 2e, arrowheads**) and a significant increase in the number of ER “whorls” (**Fig. 2e, see inset in *rab-3p::xbp-1s* animals, and see Fig. 2f for quantification of whorls**), which are often seen upon upregulation of UPR^ER^/ER-phagy (Lingwood et al., 2009; Schuck et al., 2009, 2014). These whorls were observed inside autophagosomes and often with lipid droplets as co-substrates (**Extended Data Fig. 2f, left, see arrowhead**), suggesting they are substrates for autophagy.

To look more broadly at lysosomal involvement in organelle turnover, we expanded our studies to a larger panel of organelles. We found that neuronal *xbp-1s* animals had (1) a significant decrease in lysosome related organelles (LROs), a subset of intestinal lysosomes which are also known as gut granules, (2) a quantifiable decrease in lipid droplets, and (3) a 3-fold increase in punctate of an ER-specific Rab GTPase, RAB-10, a proposed marker for ER tubulation (English and Voeltz, 2013), but no change in other organelles (**Extended Data Fig. 3a-c, Extended Data Table 1, and see **Extended Data Fig. 3d** for examples of non-colocalized markers**). RAB-10, in conjunction with the adaptor protein, EHBP-1 (an EH domain binding protein) and EHD2 (EH domain containing 2 or RME-1 in nematodes) is necessary for the autophagic engulfment of cellular LDs (Li et al., 2016). This complex has also been described in the *C. elegans* intestine, though its designated function was reported in recycling endosomes (Shi et al., 2010; Wang et al., 2016). Taken together, these data demonstrate that the ER remodeling and lipid depletion found in neuronal *xbp-1s* animals is tightly linked to autophagy, potentially through selective lipophagy of lipid droplets and ER-phagy.

### A lipophagy complex mediates CERM formation and lipid turnover and is required for lifespan extension of neuronal xbp-1s animals

We hypothesized that elements of the RAB-10/EHBP-1/RME-1 lipophagy complex might mediate the notable decrease in LDs seen in our neuronal *xbp-1s* animals. Consistent with this hypothesis, knockdown of *ehbp-1* was sufficient to suppress the lipid depletion in the intestine of neuronal *xbp-1s* animals, similar to *xbp-1* knockdown (Fig. 3a). When quantified, knockdown of *xbp-1* or *ehbp-1* in neuronal *xbp-1s* animals most effectively suppressed LD depletion (Fig. 3b-c), with RNAi knockdown of *rme-1* only partially suppressing this phenotype and knockdown of *rab-10* having little, to no, effect. This effect could be due in part to the fact that *rab-10* knockdown has been associated with induction of autophagy, which could confound a lipid rescue (Hansen et al., 2008). Moreover, knockdown of *rab-10* and *rme-1* cause phenotypes associated with calorie restriction, including egg-laying and growth defects. In contrast, animals with *ehbp-1* knockdown are largely wild-type. Taken together, these data suggest that EHBP-1, potentially through the RAB-10/RME-1/EHBP-1 complex, is involved in the ER remodeling and lipid depletion found in neuronal *xbp-1s* animals.

**Figure 3.**
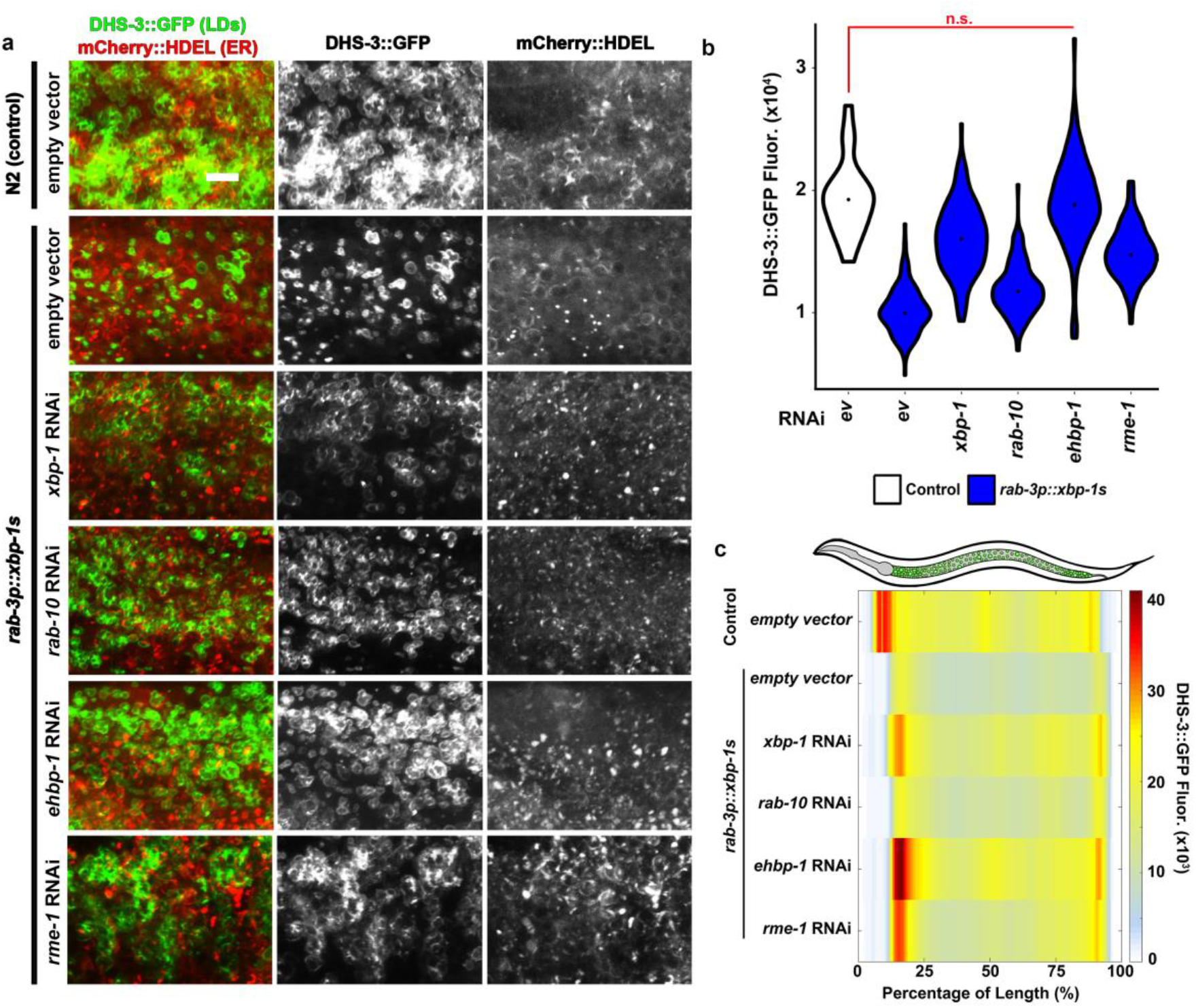
Depletion of intestinal lipids in neuronal *xbp-1s* animals is dependent on lipophagy. **(a)** Representative confocal micrographs of intestine from staged, Day 2, adults with red fluorescently-labeled endoplasmic reticulum (mCherry::HDEL) and green fluorescently-labeled lipid droplets (DHS-3::GFP) following various RNAi treatments. Scale = 5 μm. **(b)** Quantification of lipid droplets (LDs) (via DHS-3::GFP fluorescence) after various RNAi treatments. \*\*\**P*<0.001 by Mann-Whitney. n.s. = not significant. **(c)** Median profiles of DHS-3::GFP fluorescence (LDs) in **b** following alignment. All animals are positioned with the anterior side to the left. Plots are representative of three biological replicates and derived from *n* ≥ 50 organisms for each condition.

Since changes in metabolism, including ER remodeling and lipid depletion, are downstream of UPR^ER^ induction, we sought to determine whether RAB-10, RME-1, and EHBP-1 functioned upstream or downstream of peripheral UPR^ER^. To our surprise, RNAi-knockdown of any of these lipophagy components did not affect UPR^ER^ induction (Fig. 4a-c). However, knockdown of *ehbp-1* was sufficient to suppress the lifespan extension found in neuronal *xbp-1s* animals (Fig. 4d). The lifespan suppression found in neuronal *xbp-1s* animals by *ehbp-1* knockdown is not due to the potential toxicity of reducing *ehbp-1* expression as RNAi knockdown of *ehbp-1* did not shorten the lifespan of wild type animals (**Extended Data Fig. 4 and Extended Data Table 2**). Taken together, these data identify EHBP-1 as the critical component for this metabolic shift, which contributes to the lifespan extension found in neuronal *xbp-1s* animals. Perhaps more interestingly, the role of EHBP-1 in longevity does not require the distal activation of canonical UPR^ER^ (i.e. chaperone network) in peripheral tissue, as knockdown of *ehbp-1* suppresses the lifespan extension, but not the induction of peripheral *hsp-4p::GFP* in neuronal *xbp-1s* animals.

**Figure 4.**
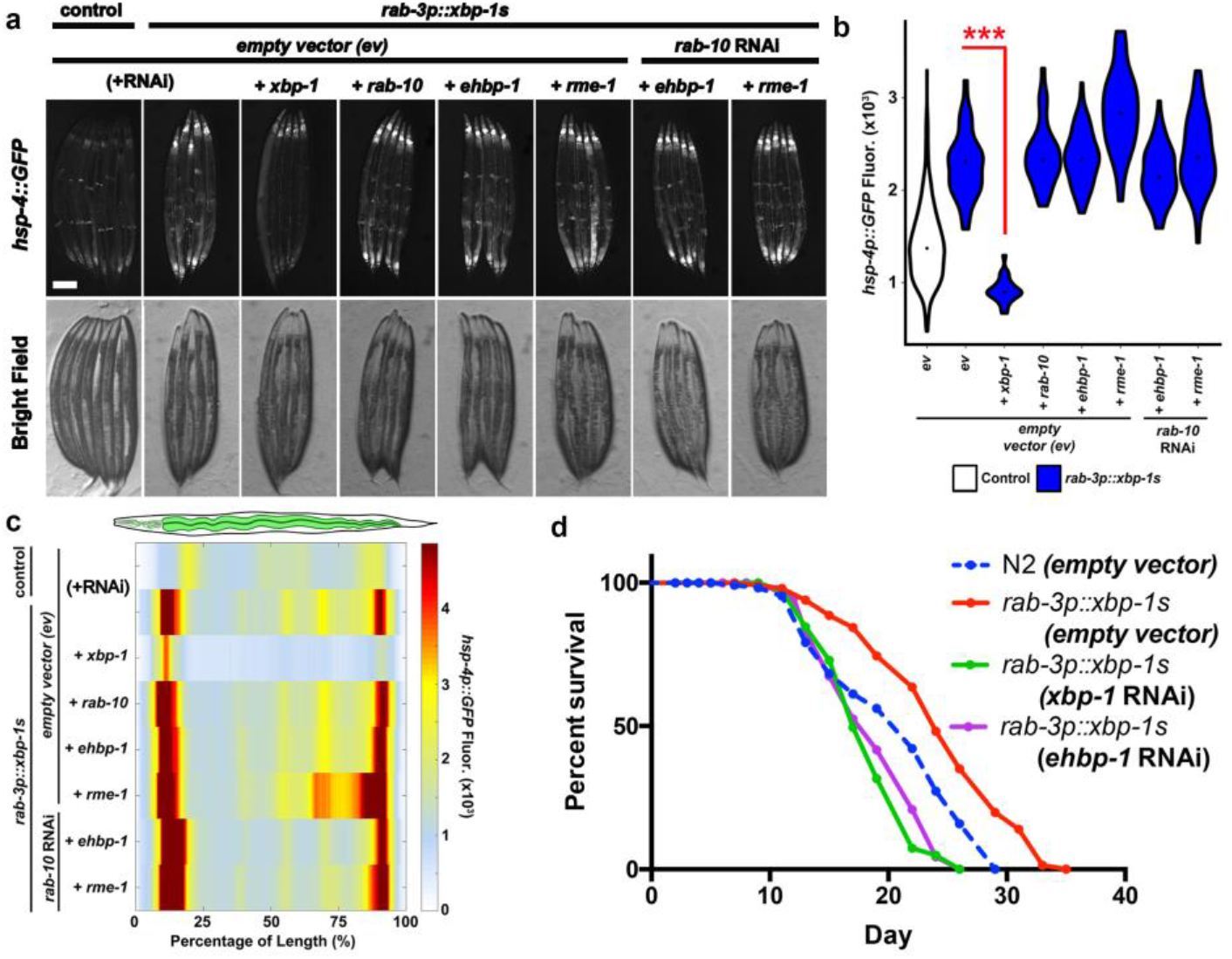
Knockdown of lipophagic components decreases longevity in neuronal *xbp-1s* animals. **(a)** Fluorescent light micrographs of *hsp-4p::GFP* UPR^ER^ reporter animals, imaged as Day 2 adults, and treated with the indicated RNAi(s) from hatch. Scale = 75 μm. **(b)** Quantification of *hsp-4p::GFP* fluorescence in control (white) and *rab-3p::xbp-1s* (blue) nematodes, using a Biosorter, following various RNAi treatments. \*\*\**P*<0.001 by Mann-Whitney. n.s. = not significant. **(c)** Median profiles of GFP fluorescence of the *hsp-4p::GFP* animals, under various RNAi treatments, in a *rab-3p::xbp-1s* background, following alignment. Plots are representative of three biological replicates and derived from *n* ≥ 50 organisms for each condition. **(d)** RNAi mediated knockdown of *ehbp-1* is sufficient to eliminate the lifespan extension normally seen in *rab-3p::xbp-1s* animals. See **Extended Data Table 2** for lifespan statistics.

Since modulation of autophagy has been found to be essential in many long-lived paradigms, we questioned whether *ehbp-1*-mediated lipophagy and ER remodeling was a critical contributor of lifespan extension in other long-lived models (Kumsta et al., 2017). Nematodes with mutations in the insulin receptor, *daf-2*, live significantly longer than wild type controls (Kenyon et al., 1993). Unlike neuronal *xbp-1s* animals, which have lower intestinal lipid content, *daf-2* animals have higher intestinal lipids (**Extended Data Fig. 5a-b**) (Vrablik et al., 2015). Moreover, we did not observe CERM formation in the intestine of these animals, and *ehbp-1* was not essential for the lifespan extension found in *daf-2* long-lived animals (**Fig. 5a, Extended Data Fig. 5b**).

**Figure 5.**
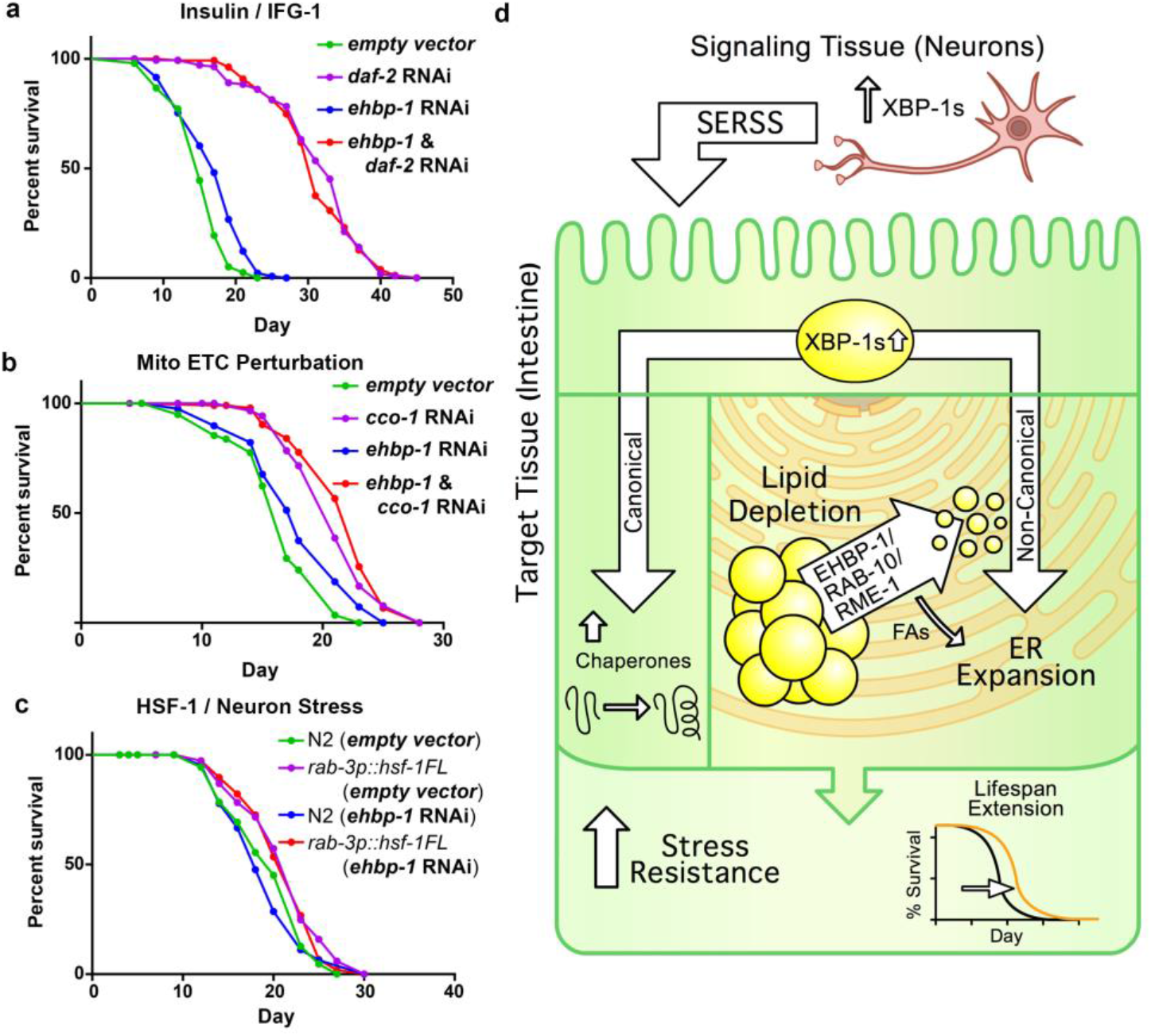
Inhibition of lipophagy does not suppress lifespan extension in other longevity models. **(a-c)** Lifespan of nematodes under normal and longevity-inducing conditions. None of these longevity paradigms is dependent on EHBP-1-mediated lipophagy. See **Extended Data Table 2** for lifespan statistics. **(d)** Controlled induction of the *xbp-1s* in neurons leads to a UPR^ER^-dependent restructuring of the ER morphology in target cells (i.e. intestine). This restructuring coincides with the lipophagic depletion of intestinal lipid droplets (LDs) and is necessary for the metabolic change and longevity seen in this model. Moreover, loss of lipophagy reverses the ER restructuring, lipid depletion, and lifespan extension in this paradigm. Finally, the longevity conferred by EHBP-1 is independent of the longevity-inducing effects of canonical chaperone induction by XBP-1s.

Mitochondrial electron transport chain (ETC) perturbation via the downregulation of a complex IV subunit *(cco-1)* has been extensively described to dramatically increase longevity (Higuchi-Sanabria et al., 2018a; Moehle et al., 2018). Similar to neuronal *xbp-1s* animals, lipids are significantly decreased in *cco-1* RNAi animals (**Extended Data Fig. 5c,e**). However, in contrast to neuronal *xbp-1s* animals, *cco-1* RNAi animals do not have CERMs and knockdown of *ehbp-1* is not sufficient to restore intestinal lipid droplets to wild type levels. Moreover, knockdown of *ehbp-1* does not suppress the lifespan extension of these animals, suggesting that lipid depletion and lifespan extension found in ETC perturbations is distinct from those identified in neuronal *xbp-1s* animals (**Fig. 5b**).

Finally, neuron-specific upregulation of the conserved heat shock transcription factor-1, HSF-1, a central player in protecting proteins from environmental and age-related stressors, has been shown to increase lifespan and stress resistance (Baird et al., 2014; Douglas et al., 2015; Higuchi-Sanabria et al., 2018b). Similar to ETC perturbation, we find that neuronal *hsf-1* animals have a significant decrease in lipid droplets compared to wild type animals. Additionally, these animals do not have CERMs and the lipid depletion is independent of *ehbp-1* as RNAi knockdown of *ehbp-1* does not rescue lipid droplet content in these animals (**Extended Data Fig. 5d-e**). Moreover, knockdown of *ehbp-1* does not suppress the lifespan extension found in neuronal *hsf-1* animals (**Fig. 5c**). Taken together, these studies have highlighted that tissue-specific regulation of lipophagy by EHBP-1 is an integral mechanism specific to the way in which cell non-autonomous UPR^ER^ mediates longevity.

## DISCUSSION

Our lab and others have developed cell non-autonomous signaling models (i.e. neuron to intestine) to provide new paradigms in the study of longevity and organismal adaptation to stress. Previously, we developed a model in which neuronal *xbp-1s* mediates increased lifespan and ER stress resistance via a neuron-specific, secreted ER stress signal (SERSS). By characterizing the intestine, the target of this signal, we uncovered a cytosolic structure, which we termed “circular <u>ER</u>-derived <u>m</u>embranes” (CERMs), which appear integral to the metabolic restructuring of this target tissue. High magnification images revealed that the intestinal ER in this model phenotypically resembles a detoxification response, which requires increased membrane demand and activation of *xbp-1.* Intriguingly, we have observed a chaperone-independent expansion of the rough ER and autophagic destruction of lipid stores (**Fig. 5d**). Taken together, these findings highlight a novel mechanism by which fundamental cell biological changes in the intestine can enact dramatic, whole-body metabolic restructuring to mediate longevity and proteostatic health.

When we characterized the intestinal ER morphology of neuronal *xbp-1s* adult animals, we found that our ER marker, *mCherry::HDEL,* labeled very large, ring-like structures, which we named CERMs. An assessment of *C. elegans* intestinal biology found that these structures most closely resemble “late endosomes,” which label positively with the small Rab GTPase, RAB-7, a protein implicated in mediating the association of multi-vesicular bodies and lysosomes with lipid droplets (LDs) (Chen et al., 2006; Schulze et al., 2017; Wang et al., 2016). Although we were unable to colocalize RAB-7 with CERMs (data not shown), we observed strong colocalization with another Rab GTPase, RAB-10. While both of these proteins are represented in LD proteomes across many phyla, including *C. elegans,* we were more interested in the role of these proteins in LD breakdown (Brasaemle et al., 2004; Cho et al., 2007; Liu et al., 2004; Zhang et al., 2011, 2012). While RAB-10 has been shown to mediate endocytic recycling in the intestine, it also works in complex with the adaptor protein, EHBP-1, and endocytosis mediator, EMD2 (RME-1 in *C. elegans),* in lipophagy (Chen et al., 2006; Li et al., 2016; Shi et al., 2010; Wang et al., 2016). These proteins have also been associated with LD size regulation (RAB-10) and cholesterol homeostasis in the liver (EHBP1, EMD2) (Denis et al., 2011; Guo et al., 2008; Morén et al., 2012). Collectively, our results suggest that upregulation of lipophagy in the intestine is an integral part of the adaptive stress response seen in neuronal *xbp-1s* animals.

Autophagy has been observed to be upregulated in response to ER stress (Kroemer et al., 2010; Kyriakakis et al., 2017; Yorimitsu et al., 2006). Specifically, induction of ER stress, either through phenobarbital-treatment, Tm-treatment, or induction of unstable dimers of membrane proteins in the ER, are all associated with a concomitant ER expansion and induction of autophagy (Bolender and Weibel, 1973; Borgese et al., 2006; Feldman et al., 1981; Lingwood et al., 2009; Schuck et al., 2009, 2014). Across many different cell types, this ER expansion often requires the transcription factor, *xbp-1s,* its downstream target *hsp-4 (BiP/GRP78),* and/or its upstream ER-localized effector *ire-1* (van Anken et al., 2003; Iwakoshi et al., 2007; Jo et al., 2009; Lee et al., 2005; Osorio et al., 2014).

To accommodate for this ER expansion, the adaptive UPR^ER^ stress response also massively upregulates gene expression of membrane lipid enzymes (e.g phosphatidyl-choline (PC) synthesis). Numerous studies have reported direct links between the activity of choline-cytidyl transferase (CCT), a rate-limiting enzyme in PC synthesis, and ER stress. Low or inhibited CCT can induce UPR^ER^, activate *xbp-1,* and correspondingly increase CCT expression; these conditions create more membrane phospholipids to be integrated into the rough ER (Hou et al., 2014; Promlek et al., 2011; Sriburi et al., 2004, 2007). A similar response can also be observed when other lipids, such as inositol or triglyceride, are depleted, or when membrane lipid saturation is increased (Chitraju et al., 2017; Promlek et al., 2011; Volmer et al., 2013). These responses to lipid disequilibrium are interesting because they are relatively slow to activate IRE-1 compared to an acute Tm-like response. This is in accordance with our neuronal *xbp-1s* model where we do not see an appreciable increase in the UPR^ER^ reporter, *hsp-4p::GFP,* or the presence of CERMs until Day 2 of adulthood, although neuronal *xbp-1s* has been expressed since birth of the animal. Also in accordance with our data, secretion is not impaired upon lipid disequilibrium or induction of *xbp-1,* but rather, it is increased following ER membrane expansion (van Anken et al., 2003; Iwakoshi et al., 2007; Lee et al., 2005; Osorio et al., 2014; Schuck et al., 2009).

The mechanism(s) by which neuronal *xbp-1s* cell non-autonomously promotes longevity and organismal stress resistance are currently unknown. Previous observations showed that neuronal *xbp-1s* increased the UPR^ER^ responsiveness of distal tissues in old age to protect against ER stress and increase longevity (Taylor and Dillin, 2013). From our observations, it is possible that ER expansion and lipophagy in the intestine is a major component of this phenomenon, in addition to induction of the UPR^ER^ (**Fig. 5d**). Increased ER membranes, especially rough ER, will have three major effects; (1) greater spacing of folding intermediates in the ER decrease the possibility of proteins aggregating, (2) sufficient and balanced lipids in the ER membranes prevent unintended membrane protein dimerization (to protect hydrophobic domains) and (3) a greater spacing of the ERAD and glycosylation machinery in the membrane will ensure they are working at full capacity (Volmer and Ron, 2015). These changes in lipid composition can be sensed by the IRE-1 machinery and induce UPR^ER^ independently of any unfolded protein stress in the ER lumen (Volmer et al., 2013). Notably, this ER response has been shown to work independently of ER chaperone levels, which also explains the low level of *hsp-4p::GFP* induction we see in our model relative to Tm treatment (Schuck et al., 2009). Intriguingly, we also found that blocking lipophagy (via *ehbp-1* RNAi) inhibits the lifespan extension of neuronal *xbp-1s* animals, but left distal *hsp-4* induction unaffected. It appears that canonical *hsp-4* induction can be separated from CERM formation and lipid depletion, indicating that chaperone induction is not solely sufficient to extend lifespan in our neuronal *xbp-1s* paradigm. This is particularly interesting by the fact that both chaperone induction and CERM formation requires *xbp-1* in the periphery and yet, autonomous induction of *xbp-1s* in the periphery, or acutely via Tm, is not sufficient to induce either CERMs or lipid depletion (Kyriakakis et al., 2017). Thus, an expanded ER, with the correct balance of lipid species, should protect animals from most conventional ER stressors, delay the age-onset of ER decline, and extend lifespan.

A second possible way that changes in intestinal ER might affect longevity is through hypersecretion and/or caloric restriction (CR). Notably, we did not observe changes in intestinal ER or LDs until Day 2 when the germline had reached its maximum speed of production. This suggests that the intestine might be able to cope with its secretory demands up to the point when apolipoprotein particle (containing vitellogenins for egg maturation) secretion ramps up. The hyper-secretory remodeling of the ER (“whorls”) has been observed during the detoxification response in liver and, in our model, may have a diluting effect on potential pathogens that try to colonize the gut (Bolender and Weibel, 1973; Feldman et al., 1981). Conversely, increased membrane demand needed by the ER expansion, which leads to lower intestinal lipids, might also cause CR, which would induce longevity as well. CR does not seem to be induced by neuronal *xbp-1s* since the extended lifespan of *rab-3p::xbp-1s* animals can be further extended by crossing into a *rab-10* (allele) mutant background (a published CR model) (data not shown, Hansen et al., 2008). This fits with previous data that suggests a *daf-16* dependence on neuronal *xbp-1s* longevity, but not *daf-2* (the insulin receptor in *C. elegans)* or *pha-4* (the FOXA transcription factor required for CR-mediated longevity in *C. elegans* (Kimura et al., 1997; Panowski et al., 2007; Taylor and Dillin, 2013).

Finally, although this phenotype was described in *C. elegans,* the phenomena of cell non-autonomous UPR^ER^ transmission from the brain to the liver, for instance, is a conserved and well-described process (Deng et al., 2013; Frakes and Dillin, 2017; Williams et al., 2014). It is envisioned, due to the conservation of the RAB-10/EHBP-1/RME-1 complex that these findings might also apply to understanding the health-promoting effects seen in a liver responding to signals from the nervous system under conditions of activated UPR^ER^.

### Methods

#### Strains and maintenance

All *C. elegans* strains used are derivatives of N2 from Caenorhabditis Genetics Center (CGC) and are listed in **Supplemental Table 1**. All worms were maintained at 15°C on NGM agar plates and fed the OP50 *E.coli* B strain of bacteria. To avoid genetic drift, all worms were maintained for a maximum of 4 months (16-20 generations) before a fresh thaw was performed. For experimentation, worms were bleached to synchronize by degrading carcasses with bleach solution (1.8% sodium hypochlorite, 0.375M KOH), then washing intact eggs 4x with M9 solution (22 mM KH_2_PO_4_ monobasic, 42.3 mM Na_2_HPO_4_, 85.6mM NaCl, 1 mM MgSO_4_), and plating eggs onto RNAi bacteria on NGM agar plates containing 1 μM IPTG, 100 μg/mL carbenicillin, and 10 μg/mL tetracycline until the desired stages of adulthood. RNAi bacteria are derived from the HT115 *E. coli* K12 strain of bacteria containing pL4440 empty vector control or expressing double-stranded RNA containing the sequence of the target gene. RNAi strains were all isolated from Vidal or Ahringer libraries and sequence-verified prior to use. All RNAi used in this study have been knockdown verified previously (Carrano et al., 2009; Durieux et al., 2011; Merkwirth et al., 2016; Tian et al., 2016).

*rgef-1p::xbp-1s* worms were synthesized by injecting N2 worms with pRT5 at 25 ng/μL, pEK2 *(myo-2p::tdtomato)* at 2.5 ng/μL and 100 ng/μL of pD64 vehicle as filler DNA. Worms positive for *myo-2p::tdtomato* were selected to identify for stable arrays. Integration was performed by gamma irradiation. Briefly, L4 worms were irradiated with 4000-4400 rems of radiation and integrants were identified by selecting animals that maintained *myo-2p::tdtomato* at 100% frequency in the F3 generation. Two independent lines were isolated, backcrossed to N2 animals 8x to eliminate mutations, and animals with the most similar phenotypes to the array animals were used for experimentation.

#### Sample preparation and Microscopy

*Worm Fixing Protocol and Sample Imaging for confocal microscopy:* Staged worms were rinsed off plates with M9 and added to 15-mL conical tubes. Animals were washed 3x with M9 to remove bacteria (do not exceed ~1,000g; 1,000 RCF on an Eppendorf centrifuge 5702), then transferred to microcentrifuge tubes using a glass Pasteur pipette (Note: glass is always recommended for transferring of worms since they will adhere to plastic surfaces). Animals were pelleted and excess M9 was aspirated to 100 μL, and 100 μL of buffered paraformaldehyde was added directly to worms (“PFA” 4% paraformaldehyde, 340 μM CaCl_2_, 1.2% Glucose, 40 mM NaH_2_PO_4_, 60 mM Na_2_HPO_4_, pH 7.4). Tubes were rocked at room temperature for 20 minutes, ~100 μL of liquid was removed, and 1.3 mL of M9 was added. Animals were then left at 4°C overnight. Animals were then spun down, solution was aspirated down to 100 μL, and 200 μL of 70% glycerol buffered in PBS was mixed in. Animals were stored at -20°C until being imaged.

For imaging, samples were mounted directly on glass slides, a coverslip was placed directly on the specimen, and sealed with nail polish. Mounted slides were kept at 4°C for at least two days prior to imaging to allow for settling of specimen and rehydration with glycerol. Fixed animals were viewed on a Zeiss LSM700 Inverted confocal microscope, with constant acquisition settings when comparing specimens within a given experiment. Unless otherwise noted, images are maximum intensity projections (using the ‘Processing’ tab in Zen software) of stacks spanning the width of various tissues within a single transgenic animal. Neutral lipid (BODIPY 409/503) staining was performed as previously described in Klapper et al., 2011.

#### Functional Lysosome Labeling and live-cell imaging using wide-field microscopy

Adapted from Hermann et al., 2005 – To label lysosomes in live worms, Lysotracker Green, [1 mM] in DMSO, was diluted at 1:100 in M9. This mixture was then spotted directly onto an OP50 bacterial lawn of NGM plates (100 μL of lysotraker was plated onto 6 mm plates seeded with 200 μL of OP50 bacteria) and left to dry in a light-proof box. Staged worms were plated onto these lysotracker plates and incubated at 20°C for two hours. Worms were then transferred to a clean OP50 *E.coli* B strain seeded plate, and incubated at 20°C for one hour to remove excess dye not taken up by lysosomes. To quantify whole worm Lysotracker signal, worms were washed off plates, moved to 15-mL conical tubes, and run on a Biosorter, and analyzed as previously described (Daniele et al., 2017). When preparing samples for microscopy, individual worms were picked off plates and mounted directly on microscope slides in M9 containing 100 nM sodium azide. A coverslip was placed directly on the sample, sealed with nail polish, and imaged immediately. Specimens were viewed on a Zeiss AxioObserver.Z1 microscope equipped with a lumencor sola light engine and a Zeiss axiocam 506 camera, driven by Zeiss ZenBlue software using a 63x/1.4 Plan Apochromat objective and standard GFP filter (Zeiss filter set 46 HE). All live-cell imaging for *mCherry::HDEL* or *dhs-3::GFP* were performed using similar methods on the same microscope. For mCherry imaging, a standard dsRed filter was used (Zeiss filter set 43).

#### Biosorter Analysis

Staged worms were washed off plates and resuspended in M9 into 15 ml conical tubes. We used a Union Biometrica complex object parameter analysis sorter (COPAS) Biosorter (product no. 350-5000-000) using both 561 nm and a 488 nm light sources. Biosorter calibration, cleaning, and sample running were performed as described in (Daniele et al., 2017).

#### Software Specifications and Data Analysis

Worm profile data was collected using the Biosort 5401.1 software provided for use with the biosorter machine. Prior to running an orientation algorithm (“LAMPro”, Daniele et al., 2017), all profiles were converted from one of multiple .dat file(s) into a single .txt file using the “Export as Text” function in the Profile Reader 16.1 software (also sold by Union Biometrica) and then saved in a folder alongside the .lmd and .dat files that were saved by the Biosorter at the time of data acquisition. All orientation algorithms were written in Perl5 and can be interpreted in Cygwin (a Linux API that can run on Windows) through additional Perl5 packages. Cygwin can be downloaded at the following address – Cygwin.com/install.html. During Cygwin installation users will be given the option to install the additional Perl5 packages. Data analysis, significance testing, and data plotting was run using MATLAB scripts (version R2015a) which have been integrated into a single graphical user interface (“LAMPro Suite 2.0”). To quantify the number and/or size of fluorescently labeled cell markers (e.g. MLS::GFP, LMP-1::GFP, GFP::RAB-10 punctae) stacks were normalized to the same threshold on a dark background using ImageJ. A smooth function eliminated scattered pixels. The area containing the intestine was highlighted and the ‘Analyze Particles’ function was used to give a count. These counts were normalized to “control” images (e.g. *rab-3p::xbp-1s* signal divided by the count from animals without *rab-3p::xbp-1s).* Enrichment relative to controls was then calculated for at least 3 biological replicates and a *t-test* was calculated to measure if this was above or below 1. Colocalization was calculated using the “JACoP” or (Just another Colocalization Plugin) for experiments that contained at least 3 biological replicates and a *t-test* was performed to see if the variance of the set was above or below matched controls. All violin plots were made in R.

#### Lifespan Analysis

Lifespan measurements were performed on solid NGM agar plates spotted with RNAi bacteria (HT115 *E. coli* strain K12). Worms were synchronized by bleaching, plated onto RNAi from hatch, and grown to adulthood at 20°C. Adult worms were moved away from progeny by moving worms onto fresh RNAi plates every day until D7-10 when progeny were no longer visible. Animals were then scored every 1-2 days for death until all animals were scored. Animals with bagging, vulval explosion, or other age-unrelated deaths were censored and removed from quantification. Prism5 software was used for statistical analysis. Log-rank (Mantel-Cox) method was used to determine significance.

#### Epifluorescence Microscopy

Staged worms were picked at random (under white light) from a population and immobilized in 100 nM sodium azide. Immobilized worms were aligned on a solid NGM agar plate and images were captures using a Leica M250FA automated fluorescent stereo microscope equipped with a Hamamatsu ORCA-ER camera.

#### Induction of ER stress

Protocol was adapted from Tian et al., 2016. Briefly, synchronized D2 stage animals were treated with 25ng/μl solution of tunicamycin in M9 buffer for 6 hours. Control animals were treated in an equivalent solution of DMSO in M9 buffer.

#### Electron Microscopy

Whole worm samples were processed for electron microscopy as previously reported. (McDONALD and Webb, 2011) Briefly, staged worms were subjected to high pressure freezing (Bal-Tec HPM 010) and then freeze substituted with an acetone/resin series (25% resin, then 50%, 75%, and 100%). Worms were cured in pure resin and then sectioned (70 nm sections) and then imaged (FEI Tecnai 12 Transmission EM) on formvar coated mesh grids.

## AUTHOR CONTRIBUTIONS

J.R.D. helped in developing the story, performed lifespans, completed all confocal and most Biosorter imaging and data analysis, prepared the figures, and wrote the manuscript. R.H.S. helped in developing the story, performed lifespans, completed wide-field imaging and data analysis, prepared the figures, and wrote the manuscript. V.R. performed lifespans, UPR^ER^ activation experiments, and Biosorter imaging. M.S. performed all sample preparation and imaging for electron microscopy. J.D., S.U.T., and J.W.P., S.M., and M.M. performed lifespans. D.J.E. made essential additions to the LAMPro software which enabled data analysis in this paper to be performed. A.D. helped in developing the story and assisted in manuscript preparation.

## COMPETING FINANCIAL INTERESTS

None to report.

## ACKNOWLEDGEMENTS

We are grateful to Lindsay Daniele for her figure illustrations and to Raz Bar-Ziv, Hope Henderson, Andrew Murley, and Ashley Frakes for careful reading of the manuscript and helpful suggestions.

